# β-glucan reprograms macrophages to attenuate efferocytosis of cancer cells

**DOI:** 10.1101/2024.11.11.622862

**Authors:** Alexandros Chatzis, Jakub Lukaszonek, Dimitris Lagos, Dave Boucher, Ioannis Kourtzelis

**Affiliations:** Hull York Medical School, University of York, United Kingdom; York Biomedical Research Institute, University of York, United Kingdom; Department of Biology, University of York, United Kingdom

**Keywords:** Macrophages, phagocytosis, efferocytosis, trained innate immunity, trained immunity, inflammasome, interleukin-1β, melanoma, breast cancer

## Abstract

Macrophage phagocytosis has been implicated in regulating anti-tumour immunity. Trained innate immunity (TII), induced *via* modulation of mature myeloid cells or their bone marrow progenitors, mediates sustained increased responsiveness to secondary challenges. Despite the advances in the study of TII-mediated anti-tumour activity, the impact of TII on the orchestration of phagocytosis in the tumour setting requires further elucidation. Here, we investigated whether macrophage phagocytosis of tumour cells can be modulated through induction of TII.

To this end, mice were pre-treated with β-glucan, a fungal-derived agonist of TII, and bone marrow was isolated for macrophage differentiation. Macrophages were then co-cultured with tumour cells that were either apoptotic or opsonised with an antibody recognising a tumour antigen, to mimic efferocytosis and antibody-dependent cellular phagocytosis (ADCP), respectively.

While TII did not have any impact in the modulation of ADCP, efferocytosis was decreased in trained macrophages. Along the same line, gene expression analysis demonstrated that mRNA levels of molecules promoting efferocytosis were downregulated in trained macrophages. Trained macrophages exerted decreased levels of active caspase-1 and produced decreased levels of interleukin-1β upon efferocytosis of tumour cells.

Our findings reveal a hitherto unknown role of TII in the regulation of anti-tumour immunity and may set the stage for designing new cancer immunotherapeutic approaches targeting macrophage efferocytosis.

## Introduction

The identification of novel mechanisms to harness immune responses is essential for bolstering tumour suppression. There is an increasing need to better understand the regulation of macrophage responses to cancer to inform the development of novel therapeutic approaches and to increase the efficacy of current immunotherapies.

Macrophages represent a major orchestrator of immune responses in a plethora of pathologies^1–3^. While residing in the tumour microenvironment, macrophages display remarkable heterogeneity and plasticity turning into essential components that can either promote or suppress tumour growth^4–6^. The tightly coordinated recognition and uptake of ‘foreign’ material through the process of phagocytosis is a major macrophage effector function that shapes host immune responses to both microbial and sterile stimuli^7,8^. In the tumour setting, macrophages can engulf cancer cells, that are opsonised with antibodies recognising surface tumour antigens, through antibody dependent-cellular phagocytosis (ADCP)^9^ and mediate the clearance of apoptotic cancer cells through the specialised type of phagocytosis called efferocytosis^10^. These two different types of phagocytosis, that represent a cardinal function of macrophages, have been linked to distinct effects on tumour growth^11^. While ADCP promotes anti-tumour effects^9^, efferocytosis promotes a tumour-tolerant condition by inducing the release of immunosuppressive mediators such as interleukin 10^10,12^. Inhibition of signaling pathways or phagocyte receptors that upregulate efferocytosis, such as Mer proto-oncogene tyrosine kinase (MerTK), has therefore been effective in reducing tumour growth in several experimental cancer models^13,14^.

Trained innate immunity (TII or trained immunity) defines long-lasting adaptations of innate immune cells based on transcriptional and epigenetic modifications of myeloid cells and their bone marrow progenitors^15–17^. Specifically, TII represents a state of enhanced immune responsiveness of the myeloid compartment to secondary challenges. We have previously shown that TII contributes to tumour suppression using experimental models of melanoma and lung cancer^18^. The anti-tumour effects of TII were attributed at least in part to epigenetic and transcriptomic reprograming in neutrophils and enhanced tumour cell killing by release of reactive oxygen species^18^. The tumour suppressive phenotype of TII has been corroborated by several studies employing different tumour models and utilising different agents to induce TII^19–22^. While the role of TII in promoting anti-tumour immunity is established, the impact of TII in efferocytosis or ADCP to shape anti-tumour immunity remains poorly explored.

The application of immunotherapies including checkpoint inhibitors that target the adaptive immune compartment for the treatment of melanoma has significantly improved the clinical management of this malignancy^23^. However, current drugs have significant limitations as some patients suffer from side effects or do not benefit from treatment^24–27^.

Macrophages are abundant in the tumour microenvironment of melanoma and their therapeutic targeting may be a promising strategy to complement existing therapies and enhance treatment success^28,29^. Of note, macrophages have been shown to contribute to ADCP^30^ of melanoma cells. In addition, targeted inhibition of efferocytosis incites anti-tumour immunity in melanoma^31^.

Here, we studied the role of TII on macrophage phagocytosis of melanoma cells and the cytokine profile of phagocytic macrophages. Our data demonstrate that ‘trained’ macrophages show decreased efferocytosis of tumour cells accompanied by reduced secretion of IL-1β. These findings unravel a hitherto unrecognized role of TII in the control of macrophage anti-tumour functions. The decreased macrophage efferocytosis of tumour cells has the potential to be harnessed therapeutically.

## Materials and Methods

### Mice and mouse experiments Mice

C57BL/6 wild type (WT) mice and Fcer1gtm1Rav (FcγR^-/-^; provided from Dr. James Hewitson, University of York) mice^32,33^ were bred in-house under specific pathogen-free conditions on a standard 12/12 h light/dark cycle according to the institutional guidelines at the Biological Services Facility (BSF), University of York. Male and female mice that were 8-12 weeks old were used in experimental procedures. Food and water were provided ad libitum.

### Mouse experiments

To induce trained innate immunity, mice were pre-treated with a single dose of 1 mg of β-glucan from *Trametes versicolor* or *Saccharomyces cerevisiae* (Invivogen) in PBS or with PBS vehicle alone (control) intraperitoneally (i.p.). Seven days after injection, bones were harvested and bone marrow was used for bone marrow-derived macrophage (BMDM) differentiation. All animals were visually inspected daily and were within accepted humane endpoints.

Induction of melanoma tumour was performed as previously described^18^. Briefly, mice were injected subcutaneously (s.c.) with 3 x 10^5^ B16-F10 cells into the right flank. To examine the role of trained immunity in tumour cell efferocytosis, mice were pre-treated with a single dose of β-glucan from *Trametes versicolor* (1 mg, Invivogen) in PBS or with PBS vehicle alone (control) i.p., followed by tumour cell injection 7 days later. Volume of palpable tumours was monitored with callipers and was calculated using the equation (length x width2) / 2. Immunofluorescence analysis was performed 19 days after tumour inoculation. Mice were excluded from experiments if pre-established exclusion criteria were fulfilled, for instance, tumour ulceration as an animal protocol-defined endpoint or, in cases that no tumour growth was observed.

### Ethics statement

All animal experiments were carried out under the authority of a UK Home Office Licence (project licence number PPL PP7424874) that obtained approval by the University of York Animal Welfare and Ethics Review Board. All procedures were performed in compliance with ARRIVE guidelines. No unexpected adverse events were recorded during this study.

### Generation of mouse bone marrow-derived macrophages (BMDMs) and neutrophils

The isolation of bone marrow and differentiation into BMDMs (hereafter mentioned as macrophages) was performed as previously described^34^. Briefly, bone marrow was flushed from femurs and tibias of mice and cells were plated and cultured in the presence of recombinant macrophage colony-stimulating factor (M-CSF; 20 ng/ml, PeproTech or Proteintech). Culture medium was replaced every two days, and after seven days differentiated BMDMs were used for further experiments. To isolate neutrophils, bone marrow cells, after erythrocyte lysis, were centrifuged in 62% Percoll gradient (Cytiva) as previously described^34^.

### Mouse cell lines

The B16F10 melanoma cell line (Creative Biogene) and the breast cancer cell lines PY8119 and PY230 (provided by Will Brackenbury, University of York) were utilised. B16F10 cells were cultured in Dulbecco’s Modified Eagle’s Medium (DMEM, Gibco) supplemented with 10% heat-inactivated fetal bovine serum (FBS, Gibco), 100 U/ml penicillin, 100 μg/ml streptomycin (Gibco) and 2 mM L-Glutamine (Gibco). PY8119 cells were cultured in F-12K Nutrient Mix medium (Gibco) supplemented with 5% FBS and 100 U/ml penicillin, 100 μg/ml streptomycin. Same medium was used for the culture of PY230 cells supplemented with 0.1% MITO+ serum extender (Corning).

### Flow cytometric analysis

Cell surface staining of MerTK on macrophages was performed in 5% FBS/PBS using 6 μg/ml of anti-MerTK (PE-conjugated, clone: DSSMMER, Invitrogen) or the IgG2a λ isotype control (PE-conjugated, clone: G013C12, BioLegend) for 25 minutes.

Levels of apoptosis were assessed using the annexin V apoptosis detection kit APC (Invitrogen) in combination with propidium iodide as previously described^34^.

Flow cytometric analysis of stained cells was performed on a CytoFLEX (Beckman Coulter) and data were analysed with CytExpert software.

### Immunofluorescence, imaging and quantitative analysis of efferocytosis in whole tumour sections

Melanoma tumours were collected from mice and frozen in the Optimal Cutting Temperature Compound (VWR Chemicals). For immunodetection studies, 5 μm frozen sections were fixed with acetone and staining with the following antibodies was performed: anti-cleaved Caspase-3 antibody (clone 5A1E, Cell Signalling) followed by an antibody conjugated with Alexa Fluor 647 (Thermo Fisher Scientific), anti-TYRP1/TRP1 (clone TA99, Bio X Cell) followed by an antibody conjugated with Alexa Fluor 555/561, BioLegend) and anti-F4/80 (clone BM8, Biolegend) followed by an antibody conjugated with Alexa Fluor 488 (Invitrogen). Control staining included the use of rabbit IgG (GeneTex), mouse IgG2A (clone: C1.18.4, Bio X Cell) and rat IgG in the case of cleaved caspase 3, TRP1 and F4/80, respectively.

Tissue sections were counterstained with 4′,6-diamidino-2-phenylindole (DAPI, Sigma-Aldrich) and mounted in Prolong Gold (Invitrogen). Images were scanned with a slide scanner (Axioscan Z1 Zeiss). Images of entire tumour sections were acquired at 20X magnification and analysed with QuPath^35^.

Automated quantification was performed after sample segmentation for DAPI^+^ cells and exclusion of necrotic regions. For quantification of macrophages that have engulfed apoptotic tumour cells, i.e. efferocytic macrophages, the number of cCasp-3^+^ TA99^+^ F4/80^+^ cells was counted and divided by the total tumour area. Imaging of the entire section was performed and a large tumour region averaging 39.6 x and 26.9 mm^2^ was assessed from control and trained mice, respectively.

### Cell treatments

Trained or non-trained macrophages were stimulated with 10 ng / ml lipopolysaccharide (LPS, Invivogen) for 16–18 h followed by RNA extraction and assessment of *Tnf* mRNA levels.

To induce apoptosis, cancer cells were treated with 1 μM of the protein kinase C inhibitor staurosporine^36^ for 16–18 h (Cell Signalling Technology). To induce apoptosis in bone marrow neutrophils, cells were cultured in HBSS containing 1% FBS for 16 – 18 h^34^.

To inhibit phagocytosis, macrophages were treated with the actin polymerization *inhibitor* Cytochalasin D (10 - 40 μΜ; Merck Group or Cayman Chemical Company) for 30 minutes, culture medium was changed and then phagocytic cargo was added.

To block signaling through the MerTK receptor, macrophages were treated with the inhibitor UNC2025 (0.1 μΜ, purity ≥ 98%; Cayman Chemical Company) for 1 hour prior to their co-culture with apoptotic cells or prior to RNA isolation.

Activity of caspase-1 was measured in macrophages after co-culture by incubating with FAM-YVAD-FMK FLICA (10 μM, Bio-Rad, as per manufacturer instructions) for 60 minutes with flow cytometry.

### Phagocytosis assays

Unstained macrophages or macrophages that were stained with either carboxyfluorescein succinimidyl ester (CFSE, 0.75 μM; Biosciences) or with DiD (3 μM; Thermo Fisher Scientific) were co-cultured with (i) apoptotic cells pre-labelled with 5 μM pHrodo Red SE (Invitrogen) in a 1:5 ratio of macrophages to apoptotic cells, (ii) pHrodo-labelled melanoma cells in the presence of 0.5 μg/ml anti-mouse/human Ab recognising TYRP1/TRP1 (gp75) on melanoma cells (clone: TA99, Bio X Cell) or the mouse IgG2a isotype control (clone: C1.18.4, Bio X Cell) in a 1:10 ratio of macrophages to tumour cells or (iii) pHrodo green *E. coli* Bioparticles (50 μg/ml, Invitrogen).

Flat bottom 96-well plates that were either non - TC-treated or TC-treated were used for flow cytometry and live cell imaging, respectively. Co-culture plates were spun for 1 minute at 250g prior to the initiation of the phagocytosis assay. Three technical replicates were used for each sample. For live cell imaging, the Livecyte instrument (Phasefocus) with a 10x magnification and a frame size of 1 mm x 1mm was used. At least 50 macrophages per region of interest were evaluated.

### Quantitative RT-PCR

Total RNA was extracted from cultured cells using TRIzol (Ambion Inc) according to manufacturer’s instructions and was quantified by spectrometry at 260 and 280 nm using a Nanodrop instrument (ThermoFischer). Complementary DNA was synthesized using the iScript cDNA Synthesis Kit (Bio-Rad). Real-time PCR was performed using the Fast SYBR Green Master Mix (Applied Biosystems) and gene-specific primers in a QuantStudio 3 Real time PCR detection system (Thermo Fisher Scientific). *18S* was used as an internal control for normalization. Data were analyzed using the comparative (ΔΔCt) method. The following primers were used: *18S*-F (GTT CCG ACC ATA AAC GAT GCC), *18S*-R (TGG TGC CCT TCC GTC AAT), *Tnf*-F (AAG CCT GTA GCC CAC GTC GTA), *Tnf*-R (GGC ACC ACT AGT TGG TTG TCT TTG), *Lxra*-F (CTC AAT GCC TGA TGT TTC TCC T), L*xra*-R (TCC AAC CCT ATC CCT AAA GCA A), *Abca1*-F (GGT TTG GAG ATG GTT ATA CAA TAG TTG T), *Abca1*-R (CCC GGA AAC GCA AGT CC), *Axl*-F (ATG GCC GAC ATT GCC AGT G), *Axl*-R (CGG TAG TAA TCC CCG TTG TAG A), *Gas6*-F (TGC TGG CTT CCG AGT CTT C), *Gas6*-R (CGG GGT CGT TCT CGA ACA C).

### Enzyme-linked immunosorbent assays (ELISAs)

Mouse IL-10 and IL-1β proteins were measured in cell culture supernatants using kits from Biolegend and Invitrogen, respectively, according to the instructions of the manufacturer.

### Statistical analysis

Statistical analyses were carried out with GraphPad Prism 10 software. For infection experiments, researchers were blinded to mouse genotype. This also applies to histology and flow cytometry analyses. Results are presented as mean plus standard error of the mean. Parametric and non-parametric tests were used as appropriate after testing for normality. Multiple-group comparisons were performed using analysis of variance (ANOVA) and the Tukey’s, Dunnett’s or Šídák’s multiple comparison tests. A *P* value of < 0.05 was considered to be statistically significant. Statistical information is provided in each figure legend.

## Results

### Efferocytosis of apoptotic tumour cells is decreased by trained macrophages

Having previously identified a tumour suppressive role for TII that is mediated by myeloid cells^18^, we sought to determine the impact of TII as a regulator of macrophage uptake of apoptotic tumour cells. To mimic efferocytosis conditions, macrophages pre-stained with CFSE were co-cultured with pHrodo red-labelled apoptotic melanoma cells (**supplementary Fig. 1A**). Pre-treatment of macrophages with the actin polymerization *inhibitor cytochalasin* D prior to their co-culture with tumour cells resulted in dose-dependent blockade of efferocytosis confirming the specificity of the assay (**supplementary Fig. 1B, C**). The effect of TII on macrophage efferocytosis of tumour cells was investigated using bone marrow derived macrophages (BMDMs) from the bone marrow of mice treated with β-glucan (hereafter described as trained macrophages). The increased levels of *Tnf* in trained macrophages upon their stimulation with lipopolysaccharide (LPS) confirmed the induction of TII^37^ in our experimental setting (**supplementary Fig. 2**). Flow cytometric analysis of the co-cultures revealed that efferocytosis was attenuated in trained macrophages (**Fig. 1B-D**). The dynamic behaviour of trained macrophages during efferocytosis of melanoma cells was mapped using real time imaging verifying their decreased capacity for tumour cell uptake (**Fig. 1E**).

**Figure 1.**
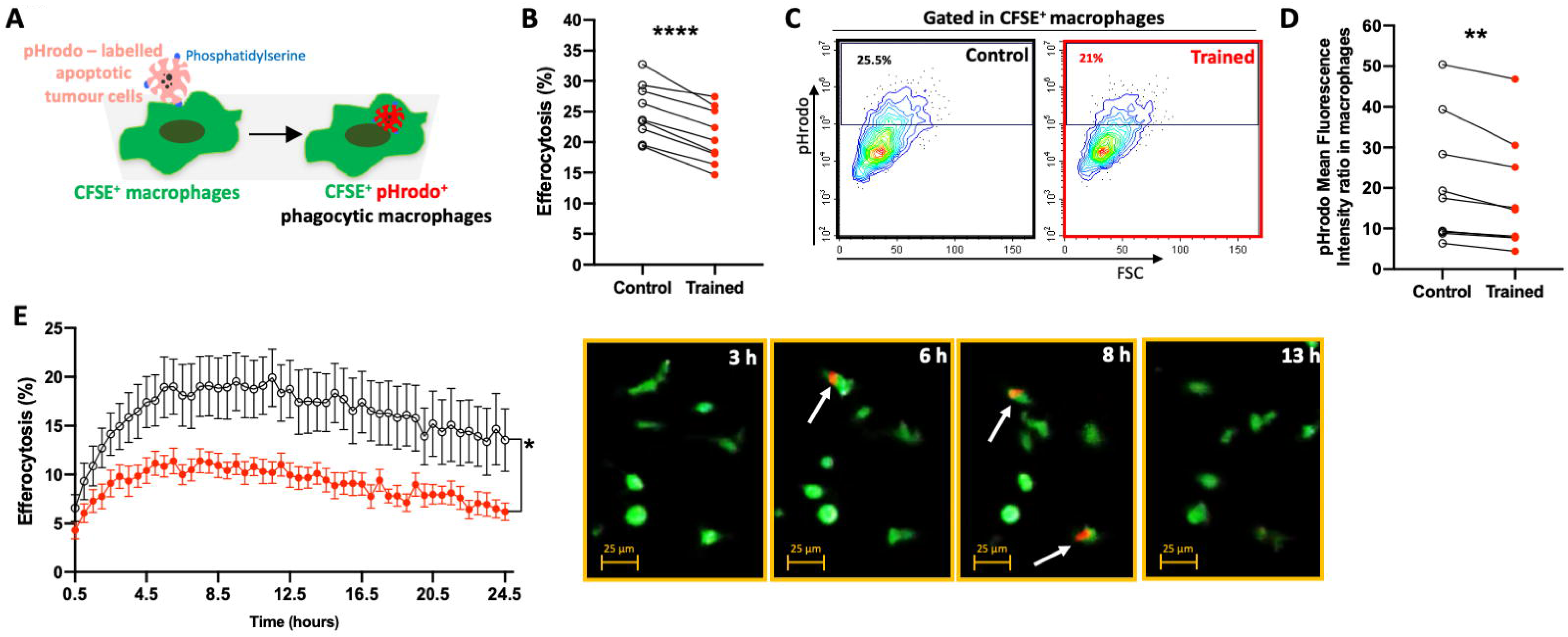
The effect of trained innate immunity on macrophage efferocytosis of melanoma cells. (A) Schematic diagram of experimental layout. (B-D) Trained or control macrophages were pre-stained with CFSE and were then co-cultured with pHrodo-labelled apoptotic B16F10 melanoma cells for 4 h. (B) Efferocytosis, (C) Representative flow cytometric plot; Numbers in outlined areas indicate the percentage of macrophages that is pHrodo^+^ red), (D) Ratio of mean fluorescence intensity (MFI) are shown and were calculated as the percentage of pHrodo^+^ macrophages and pHrodo MFI in macrophages, respectively. (B, D) Each paired line represents average values from nine different experiments (*n*LJ=LJ44 separate cell isolations per group). (E) Live cell imaging was performed in co-cultures as shown in (A). Levels of efferocytosis are shown over time (left, *n*LJ=LJ8-10 separate cell isolations per group). Representative images in selected time points are shown (right). Arrows denote phagocytic macrophages. \**P*<0.05, \*\**P*<0.01, \*\*\*\**P*<0.0001. Paired (B, D) two-tailed Student’s *t*-test, Two-way ANOVA test (E). Data are presented as mean and mean ± s.e.m.

To investigate the impact of TII in efferocytosis of tumour cells *in vivo*, we performed an immunofluorescence analysis of B16-F10 tumour sections from mice that have been treated with a single dose of β-glucan or control vehicle seven days prior to tumour cell injection as previously described^18^. The levels of macrophage efferocytosis of apoptotic melanoma cells *in situ* were significantly decreased in tumour samples from trained mice in comparison with control mice (**supplementary Fig. 3**). These data agree with our co-culture findings (**Fig. 1B-E**) further corroborating the novel role of TII on inhibition of efferocytosis in cancer.

To determine whether the effect of TII on macrophage efferocytosis applies to other types of cancer, the two breast cancer cell lines PY230 and PY8119 were used in our co-culture experiments. Consistently with our results on decreased efferocytosis of melanoma cells, we demonstrated that trained macrophages exerted attenuated efferocytic activity upon their co-culture with apoptotic breast cancer cells (**Fig. 2**) suggesting a broad role of TII on shaping efferocytosis of cancer cells.

**Figure 2.**
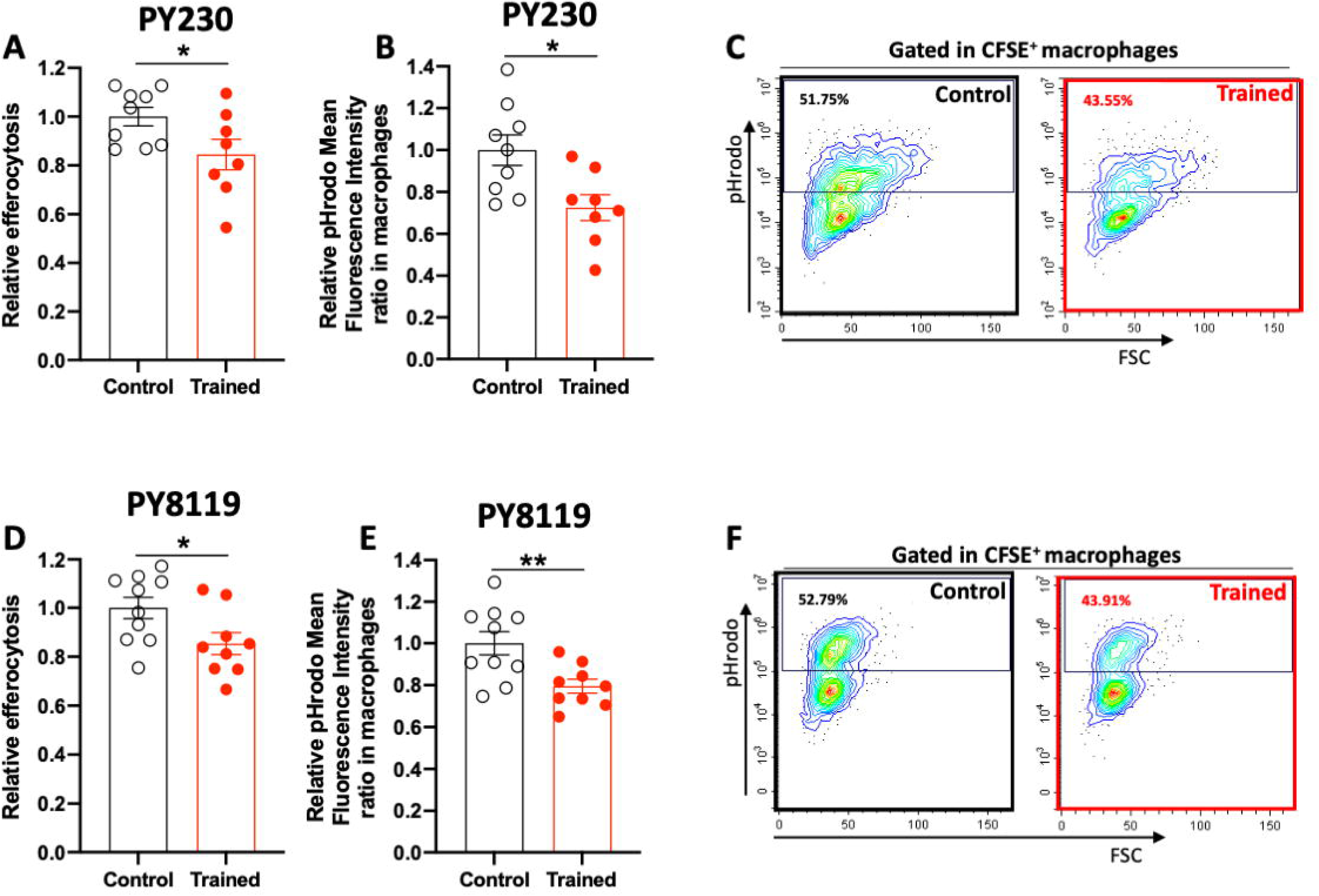
The impact of trained macrophages in efferocytosis of breast cancer cells. Trained or control macrophages were pre-stained with CFSE and were then co-cultured with pHrodo-labelled apoptotic PY230 (A-C) and PY8119 (D-F) breast cancer cells for 4 h. (A, D) Relative efferocytosis and (B, E) relative ratio of mean fluorescence intensity (MFI) are shown and were calculated as the percentage of pHrodo^+^ macrophages and pHrodo MFI in macrophages, respectively. Data are expressed relative to the control group, set as 1 (*n*=8-9 (A-C) and *n*LJ=LJ9-10 (D-F) separate cell isolations per group). (C, F) Representative flow cytometric plots are shown. Numbers in outlined areas indicate the percentage of macrophages that is pHrodo^+^ red. \**P*<0.05, \*\**P*<0.01, two-tailed Student’s *t*-test (A-F). Data are presented as mean ± s.e.m. and are pooled from two experiments (A-F).

### TII does not have any effect on macrophage ADCP of melanoma cells

We next investigated the role of TII on the macrophage ADCP. To induce ADCP, the co-culture was performed in the presence of the well-characterised anti-mouse/human TA99 Ab that recognises the tumour antigen Tyrp1^38^ (**Fig. 3A**). Specificity of ADCP induction was demonstrated using macrophages from mice deficient in Fcγ receptors^32,33^ (**Fig. 3B**). Following co-culture of TA99-opsonised melanoma cells with either trained or non-trained macrophages, levels of ADCP were comparable between the two groups (**Fig. 3C, D**), suggesting that TII does not play any role in macrophage ADCP of melanoma cells under these conditions.

**Figure 3.**
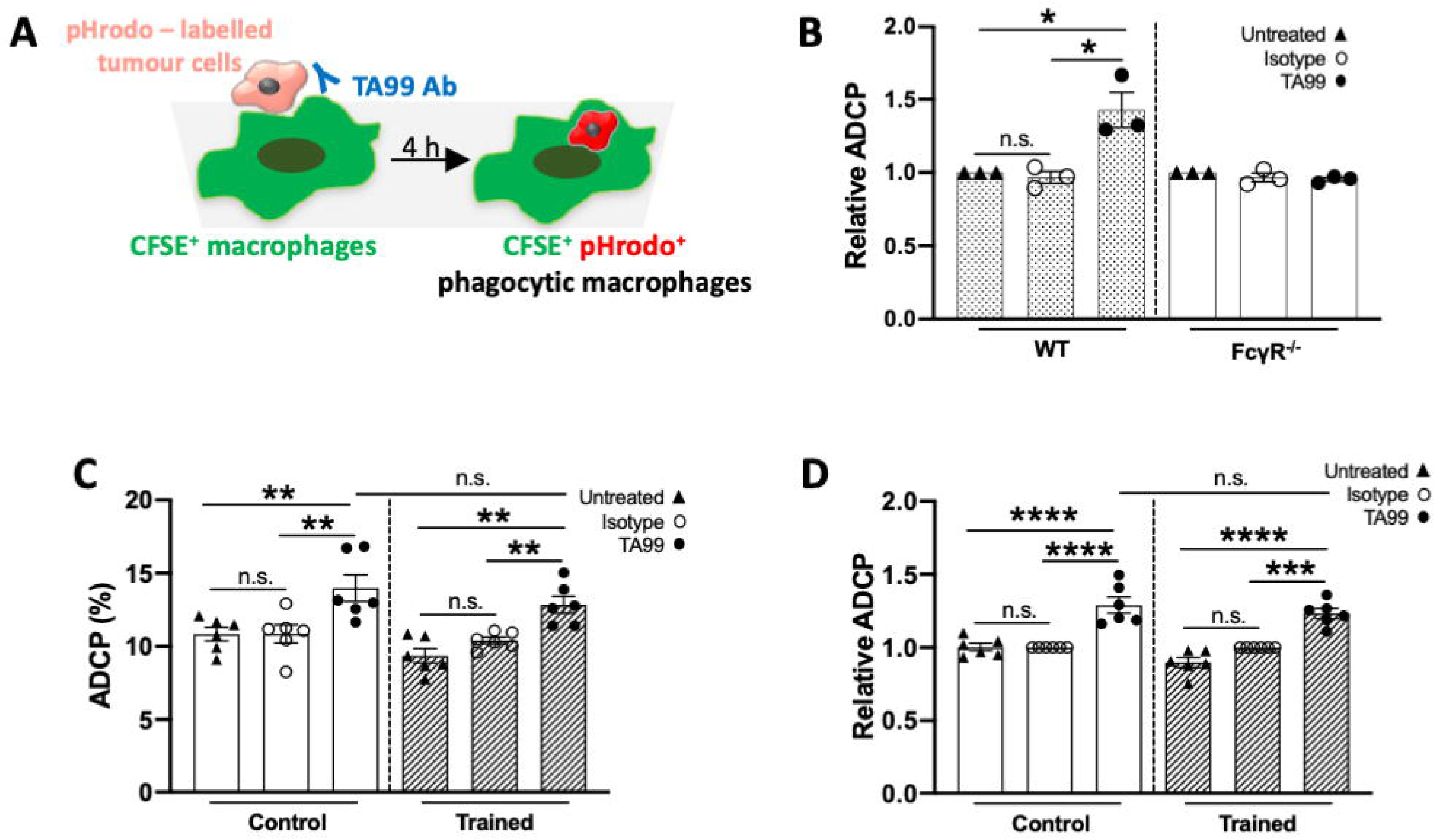
Trained innate immunity does not impact macrophage ADCP of melanoma cells. (A) Schematic diagram of experimental layout. (B) Macrophages derived from WT or FcγR^-/-^ mice were pre-stained with CFSE and were then co-cultured with pHrodo-labelled B16F10 cells in the presence of 0.5 μg/ml TA99 Ab that recognises the tumour antigen tyrosine related protein-1 (TYRP1 or gp75) or isotype control for 4 h. Relative ADCP is shown and is calculated as the percentage of pHrodo^+^ macrophages. Data are expressed relative to the untreated control group, set as 1 (*n*LJ=LJ3 separate cell isolations per group). (C, D) Trained or control macrophages were pre-stained with CFSE and were then co-cultured with pHrodo-labelled B16F10 cells as in (B). (C) ADCP and (D) relative ADCP are shown and calculated as the percentage of pHrodo^+^ macrophages. Data are expressed relative to the control group, set as 1 (*n*LJ=LJ6 separate cell isolations per group). \**P*<0.05, \*\**P*<0.01, \*\*\**P*<0.001, \*\*\*\**P*<0.0001, n.s., non-significant. One-way ANOVA with Tukey’s multiple comparisons test. Data are presented as mean ± s.e.m. and are pooled from three independent experiments (B-D).

Given that macrophage phagocytosis can be differentially modulated by different types of phagocytic cargo^39,40^, we sought to investigate the behaviour of trained macrophages towards different key phagocytic cargos. To this end, trained or non-trained macrophages were cultured together with microbial bioparticles or apoptotic neutrophils to mimic phagocytosis during infection or resolution phase of sterile inflammatory responses, respectively. The uptake of both *E. coli* particles (**supplementary Fig. 4A-D**) and apoptotic neutrophils (**supplementary Fig. 4E, F**) was elevated in trained macrophages compared to the phagocytic activity shown in control macrophages. These findings underline the ability of trained macrophages to shape differential responses to distinct phagocytic cues.

### Induction of TII combined with MerTK inhibition further downregulate efferocytosis of tumour cells

MerTK functions as an efferocytic receptor recognising the ‘eat-me’ signal PS on the outer surface membrane of apoptotic cells with the help of bridging molecules such as Gas6. As MerTK blockade has already been shown to promote tumour suppression^31^, we investigated the consequences of MerTK inhibition on trained macrophage efferocytosis.

To this end, trained or control macrophages were cultured with apoptotic melanoma cells in the presence of the small-molecule MerTK inhibitor UNC2025^41^. In trained macrophages, the presence of UNC2025 resulted in decreased efferocytosis compared to that one in control macrophages treated with the inhibitor (**Fig. 4A**). Gene expression analysis of molecules downstream of MerTK signaling^14^ in macrophages treated with the UNC2025 verified the downregulation of *Liver X receptor alpha* (*Lxra*) and *ATP-binding cassette transporter* (*Abca1*) (**supplementary Fig. 5**). We then assessed the impact of TII in the expression levels of molecules implicated in the MerTK pathway and found that the components of MerTK-dependent signaling *Gas6*, *Lxra* and *Abca1* were downregulated in trained macrophages (**Fig. 4B, C**). MerTK protein levels were downregulated in trained macrophages (**Fig. 4D**). Therefore, decreased efferocytosis of tumour cells by trained macrophages may be attributed at least in part to the downregulated molecular machinery orchestrating MerTK-dependent efferocytosis.

**Figure 4.**
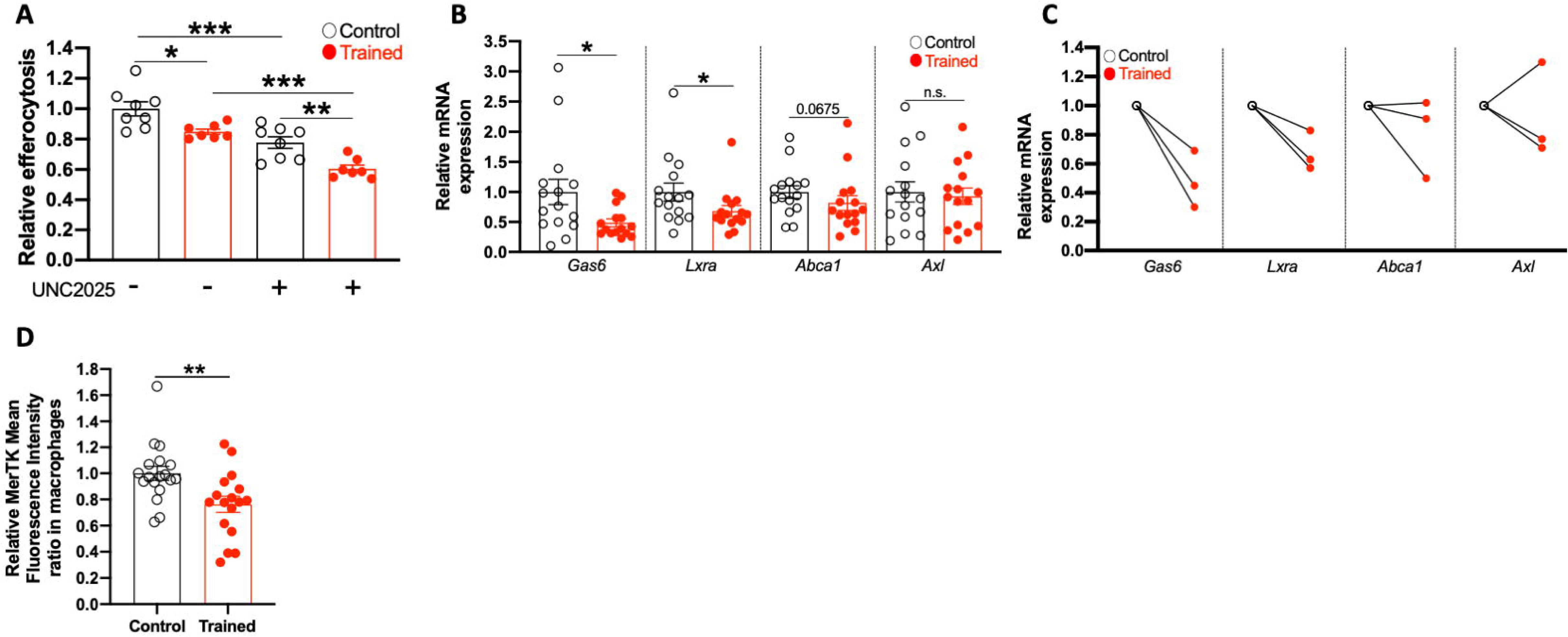
The role of MerTK pathway on the phenotype of trained macrophages. (A) Trained or control macrophages were pre-stained with CFSE, were treated with 0.1μM of the MerTK inhibitor UNC2025 or control vehicle for 1h and were then co-cultured with pHrodo-labelled apoptotic melanoma cells for 4 h. Relative efferocytosis using flow cytometry is shown and was calculated as the percentage of pHrodo^+^ macrophages. Data are expressed relative to the control group, set as 1 (*n*=7-8 separate cell isolations per group). (B, C) Relative mRNA expression of *Gas6*, *Lxra*, *Abca1*, *Axl* from control or trained macrophages. Relative mRNA expression was normalized against *18S* rRNA and was set as 1 in the control macrophages (n=15 separate cell isolations per group). (C) Each paired line represents average values from different experiments shown in (B). (D) The relative MerTK MFI in control or trained macrophages is shown. Relative MFI ratio was set as 1 in the control macrophages (n=17-18 separate cell isolations per group). \**P*<0.05, \*\**P*<0.01, \*\*\**P*<0.001, n.s., non-significant. One-way ANOVA with Tukey’s multiple comparisons test (A), two-tailed Student’s *t*-test (B, D). Data are presented as mean and mean ± s.e.m. and are pooled from two (A), three (B, C), four experiments (D).

### Secretion of IL-1β by trained macrophages upon efferocytosis of tumour cells is blunted

Macrophage efferocytosis not only paves the way towards the clearance of apoptotic cells, but also promotes metabolic alterations in macrophages^42–44^. At the same time, metabolic alterations in efferocytic macrophages induced by digested apoptotic material substantially regulates the cytokine secretion by efferocytic macrophages^45^. Since upregulation of interleukin-1β (IL-1β) has been linked to both enhanced efferocytosis^46^ and tumour growth^47,48^, we determined the levels of IL-1β production in our co-culture setting. Production of IL-1β was induced as a result of efferocytosis, as IL-1β was minimally detected in either macrophage or apoptotic melanoma cell monocultures (**Fig. 5A**). Trained macrophages displayed lower levels of IL-1β production as shown in co-culture supernatants compared to those from control macrophages (**Fig. 5A**), further underscoring the tumour suppressive effect of TII through inhibition of efferocytosis. Consistently, we demonstrated that levels of active caspase-1 were decreased in trained macrophages after efferocytosis of apoptotic tumour cells (**Fig. 5B, C**) thereby explaining at least in part the decreased levels of active IL-1β in co-culture supernatants.

**Figure 5.**
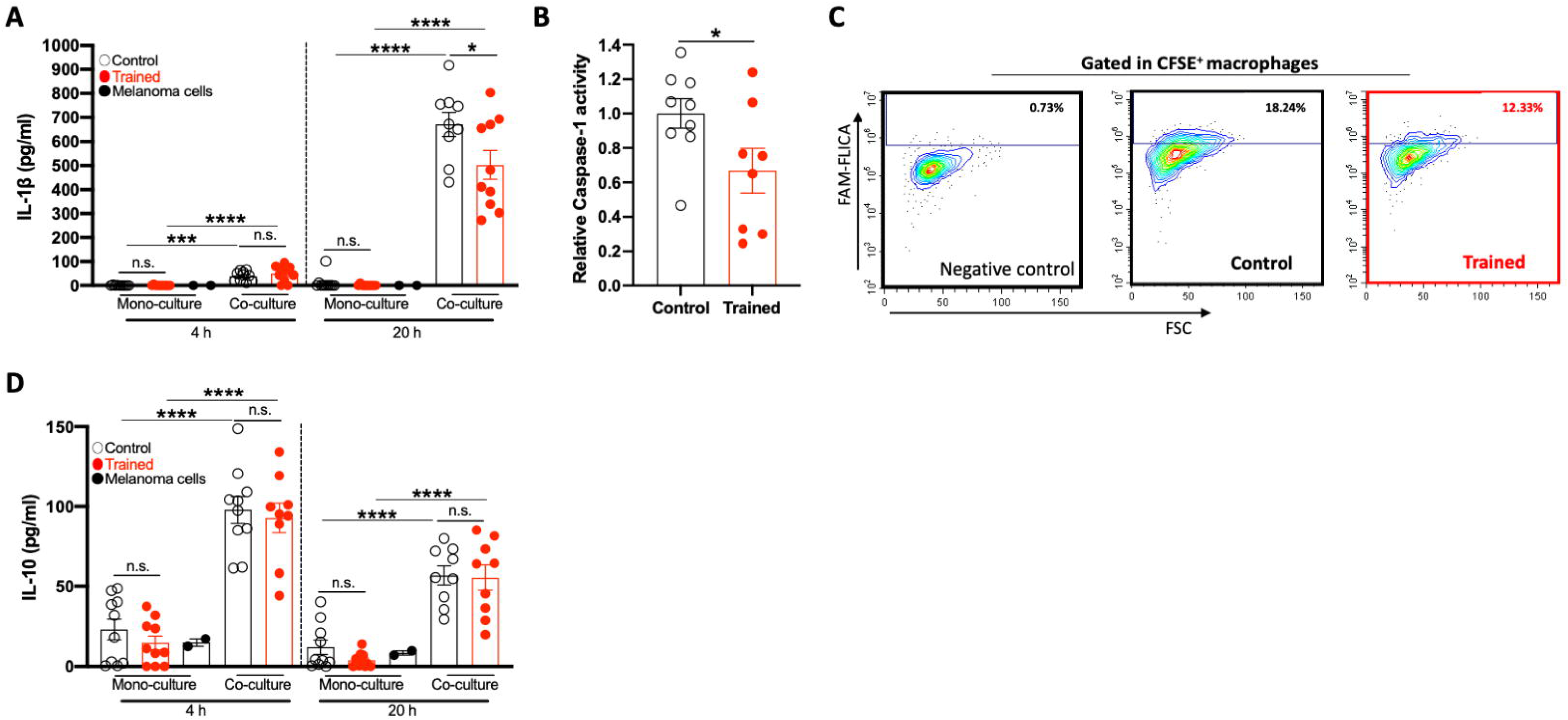
The levels of immune mediators and active caspase-1 during macrophage efferocytosis of melanoma cells. Trained or control macrophages were co-cultured with apoptotic melanoma cells for 4 or 20 h. Protein concentrations of (Α) IL-1β and (D) IL-10 were measured in efferocytosis co-culture supernatants. Monocultures of trained or control macrophages and apoptotic melanoma cells were used as controls. *N*=9-10 separate cell isolations per group for macrophages and *n*=2 for B16F10 cells. (B, C) Trained or control macrophages, pre-stained with the DiD, were co-cultured with apoptotic melanoma cells for 4 h. The probe FAM-YVAD-FMK FLICA (10 μM) was added 1 h prior to the end of the co-culture. Relative caspase-1 activity is shown and was calculated as the percentage of positive macrophages that are labelled with the FLICA (*n*=8-9 separate cell isolations per group). Data are expressed relative to the control group, set as 1. (C) Representative flow cytometric plots are shown. Numbers in outlined areas indicate the percentage of macrophages that is FAM-FLICA^+^. \**P*<0.05, \*\*\**P*<0.001, \*\*\*\**P*<0.0001, n.s., non-significant. One-way ANOVA with Šídák’s multiple comparisons test (A, D), two-tailed Student’s *t*-test (B). Data are presented as mean ± s.e.m. and are pooled from two experiments (A-D).

We also determined the levels of immunosuppressive cytokine interleukin 10 (IL-10), previously described to be upregulated after efferocytosis^12^, in the supernatants of efferocytosis co-cultures. Levels of IL-10 in the co-culture supernatant were comparable between the trained and control macrophages (**Fig. 5D**).

Together, these data unveil a novel role for TII in shaping macrophage-cancer cell interactions, providing a better understanding on mechanisms that may be targeted to block efferocytosis-dependent tumour growth.

## Discussion

The orchestration of host immune responses against cancer involves several immune cell types and events including monocyte infiltration, macrophage activation, cytokine release and phagocytosis of cancer cells, all of which aim to contribute to tumour suppression^49^. Although immune cell interactions play important roles in the immunopathology of cancer, the network of molecules and mechanisms that coordinate this complex process warrants further elucidation. Therefore, identification of immune regulatory networks that aim to harness cancer-associated immunosuppression are required. Significant advancements have been made in identifying the multifaceted roles and therapeutic potential of trained immunity in cancer^50^. In an experimental model of melanoma, induction of TII has been shown to promote myeloid cell – dependent tumour suppression through reprograming the phenotype of neutrophils to inhibit tumour growth^18^. Here, we investigated whether TII regulates macrophage phagocytosis of tumour cells. We show for first time that trained macrophages have decreased phagocytic activity of apoptotic tumour cells both in a co-culture and *in vivo* using a mouse setting. These findings were accompanied by lower levels of secreted IL-1β. We also found that *Gas6*, *Lxra*, *Abca1* and MerTK, all previously linked to enhanced efferocytosis^34,51^, were downregulated by trained macrophages. Under the tested conditions, levels of ADCP were comparable between trained and control macrophages. Use of antibodies recognising alternative melanoma surface antigens may promote ADCP by trained macrophages.

Further studies would be useful to address why trained macrophages demonstrate a cargo-dependent phagocytic behaviour, i.e. decreased efferocytosis of tumour cells, unaltered ADCP and enhanced phagocytosis of apoptotic neutrophils and *E. coli* particles. In agreement with the literature, although using a different approach to induce TII, enhanced phagocytosis of pathogen particles^37^ and apoptotic neutrophils^52^ by trained phagocytes has been demonstrated. Our findings on trained macrophage efferocytosis of different apoptotic cargos further support previous reports suggesting that the cellular identity of the engulfed apoptotic cell instruct distinct macrophage phenotypes^53^ that may be potentially linked to enhanced effectiveness of macrophage-based cell therapies.

Targeting efferocytosis to bolster anti-tumour immunity has attracted attention^10,13^. Of note, efferocytosis has been linked to cancer therapeutic resistance among several types of cancer^54,55^. The role of TII on attenuating macrophage efferocytosis in the tumour setting was corroborated by showing that inhibition of the efferocytosis receptor MerTK further decreased the uptake of apoptotic tumour cells in co-cultures with trained macrophages. As blockade of MerTK may have a direct impact in cancer cells^56^, further studies are required to address whether MerTK inhibition affects solely efferocytosis in our experimental setting.

The decreased caspase-1 activity in trained efferocytic macrophages was reflected by lower levels of IL-1β levels. Efferocytosis has been shown to induce the activation of NLRP3 inflammasome signaling in myeloid cells thereby promoting IL-1β secretion and tumour growth^46^. Further focus on the identification of specific inflammasomes that contribute to the attenuated efferocytosis by trained macrophages may support the development of novel cancer immunotherapeutic targets. In addition, future studies will provide further insight into the potential role of MerTK inhibition on macrophage IL-1β secretion during efferocytosis.

Given the macrophage heterogeneity^57^ and the different types of central and peripheral trained innate immunity^58^, the use of BMDMs as the only source of macrophages may represent a limitation of this study. However, BMDMs have been utilised extensively for the elucidation of mechanisms orchestrating cargo-dependent macrophage phagocytosis^53^. Future validation experiments using different types of macrophages will strengthen the impact of this study. In addition, utilisation of a human macrophage setting would provide additional validation of our findings. However, induction of trained immunity entails epigenetic and metabolic reprogramming of hematopoietic stem and progenitor cells (HSPCs). As such, the elucidation of mechanisms that modulate trained immunity in human cells would require the establishment of a macrophage differentiation model based on the use of HSPCs rather than the stimulation of monocytes or macrophages with β-glucan.

Our *in vivo* data are based on the use of the syngeneic B16-F10 model strengthening the impact of our co-culture findings. Future studies would be required to address further the physiological relevance of trained macrophages in efferocytosis of tumour cells using autochthonous mouse models of cancer.

Cancer combination therapies, including the administration of immunotherapy, have contributed to the improvement of the clinical management of cancer particularly in aggressive types such as melanoma^59,60^ and also breast cancer^61,62^. The anti-tumour effect of β-glucan and the vaccine Bacillus Calmette-Guerin (BCG), both inducers of TII, is being tested in clinicals trials^63–65^. Along the same line, administration of β-glucan enhances the anti-tumour immunotherapeutic effect of BCG *via* myeloid cell reprogramming^66^. TII - dependent decrease of efferocytosis may be a promising adjuvant immunotherapeutic strategy to complement existing cancer therapies and enhance treatment success.

In summary, the present study pinpoints a novel role for trained innate immunity in attenuating macrophage efferocytosis of tumour cells and demonstrates that the decreased efferocytosis by trained macrophages is associated with lower levels of active caspase-1 and IL-1β secretion. This novel role of trained innate immunity could be therapeutically exploited in pre-clinical models to expand our current mechanistic and translational understanding of the anti-tumour potential of trained immunity.

## Author contributions

AC contributed to experimental design, performed experiments, analysed and interpreted data; JL performed experiments; DL contributed to experimental design and interpreted data; DB contributed to experimental design, interpreted data and supervised research; IK conceived and designed the study, performed experiments, supervised research, interpreted data, and wrote the paper. All authors critiqued and edited the manuscript.

## Acknowledgements

We would like to thank staff at the Imaging and Cytometry Lab in the University of York Bioscience Technology. We thank Prof. Will Brackenbury (University of York) for providing the PY8119 and PY230 cell lines and Dr. James Hewitson (University of York) for providing the Fcer1gtm1Rav mice.

## Author Declarations

We confirm all relevant ethical guidelines have been followed, and any necessary ethics committee approvals have been obtained.

---

**Supplementary Figure 1.**
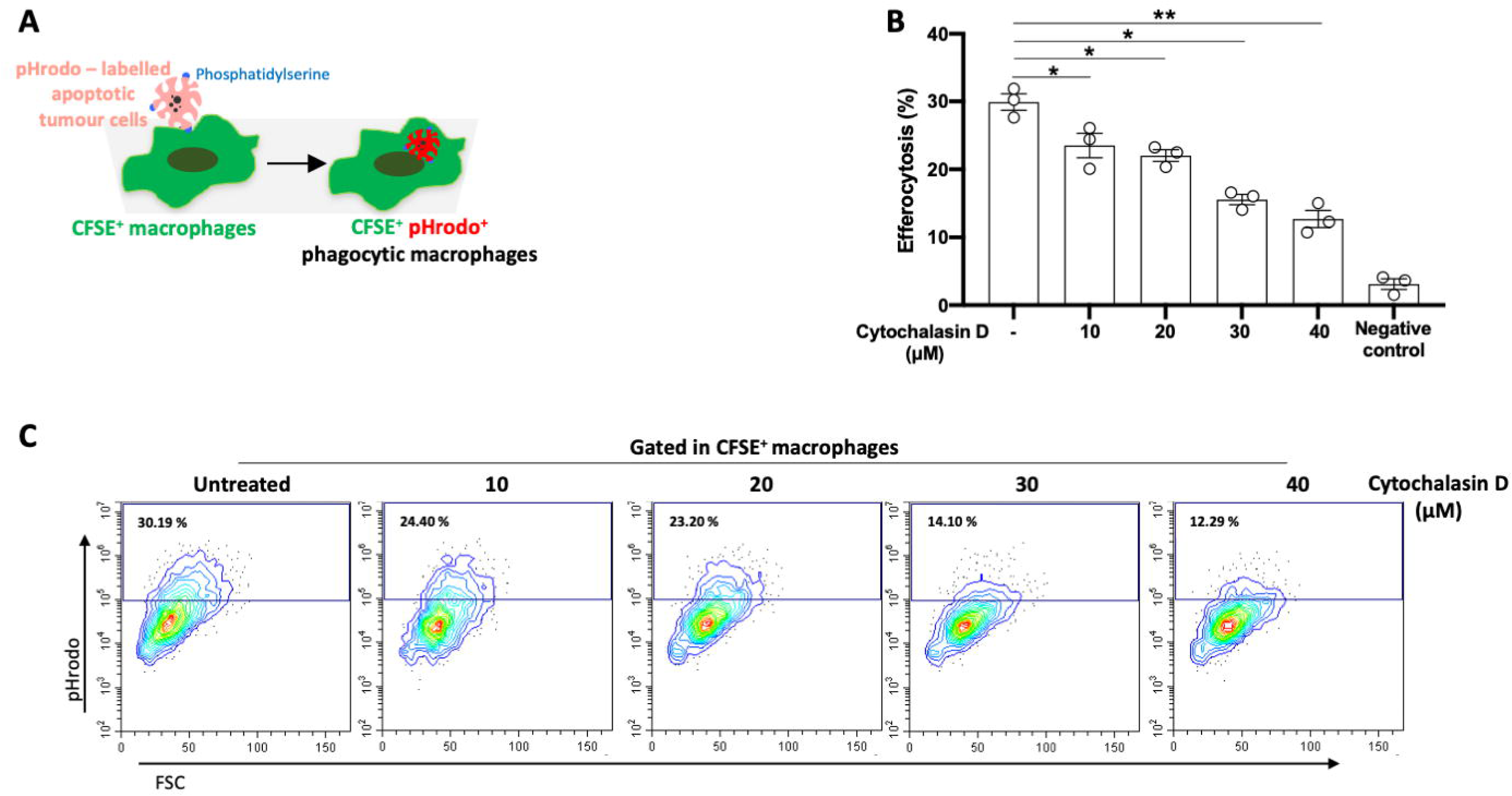
The inhibitory effect of cytochalasin D on macrophage efferocytosis. (A) Schematic diagram of experimental layout. (B) CFSE-labelled macrophages were treated with the actin polymerization *inhibitor cytochalasin* D for 30 min prior to their co-culture with pHrodo-labelled apoptotic melanoma cells for 4 h (n=3 separate cell isolations). Samples in which apoptotic cells were added to the macrophage cultures just before the flow cytometric analysis, without any incubation, served as negative control. Efferocytosis is shown as the percentage of pHrodo^+^ macrophages. (C) Representative flow cytometric plots are shown. Numbers in outlined areas indicate percentage of macrophages that is pHrodo^+^ red. \**P*<0.05, \*\**P*<0.01, One-way ANOVA with Dunnett’s multiple comparisons test. Data are presented as mean ± s.e.m. and are derived from one experiment.

**Supplementary Figure 2.**
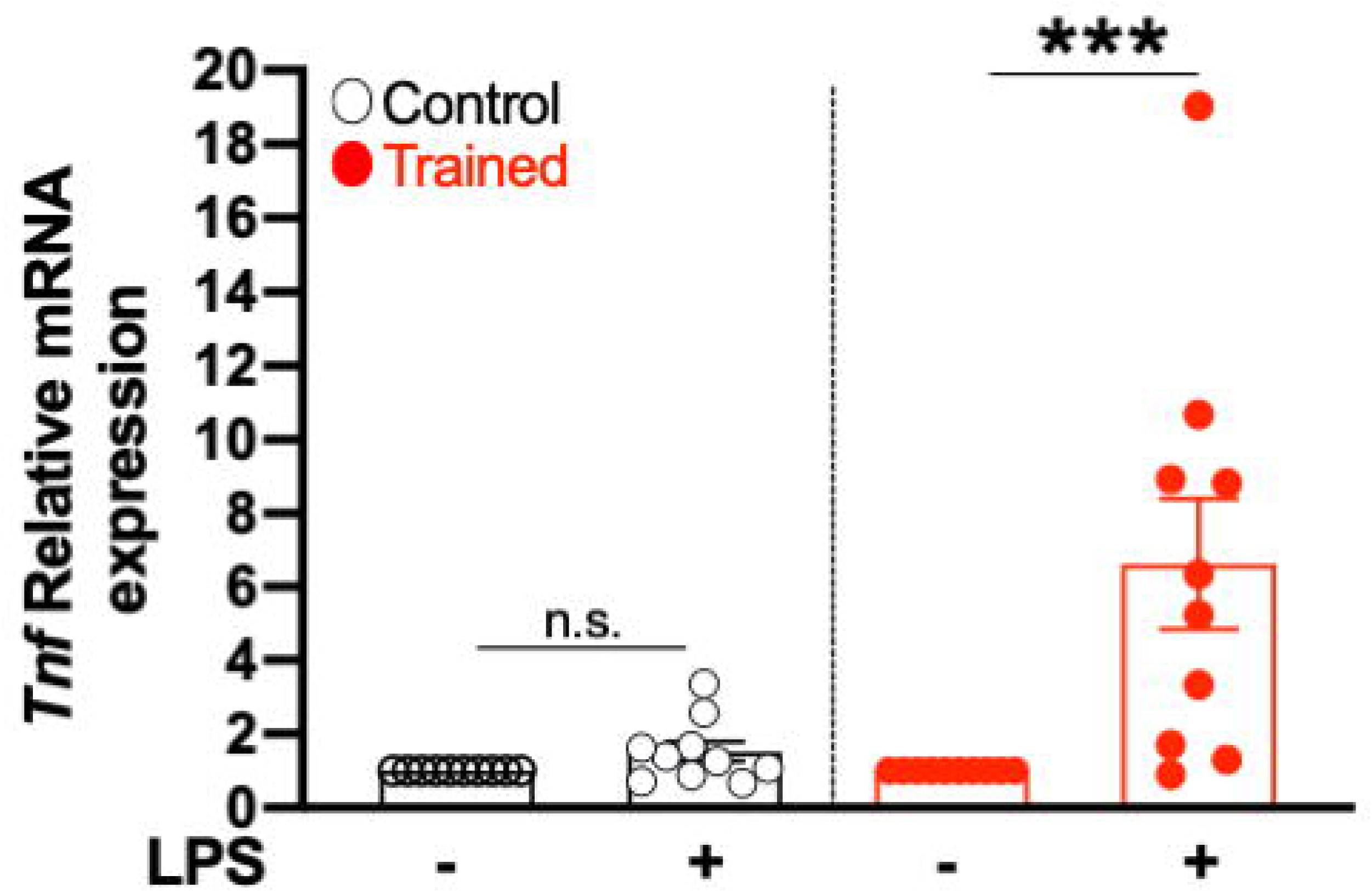
The effect of LPS on *Tnf* mRNA expression levels in trained macrophages. Relative mRNA expression of *Tnf* in trained and control macrophages that were cultured in the presence or absence of 10 ng/ml LPS for 16-18 h. Relative mRNA expression was normalized against 18S rRNA and was set as 1 in macrophages that were not treated with LPS (n=10 separate cell isolations for control and 9 for trained macrophages). \*\*\**P*<0.001, n.s., non-significant. Mann-Whitney test. Data are presented as mean ± s.e.m. and are pooled from three experiments.

**Supplementary Figure 3.**
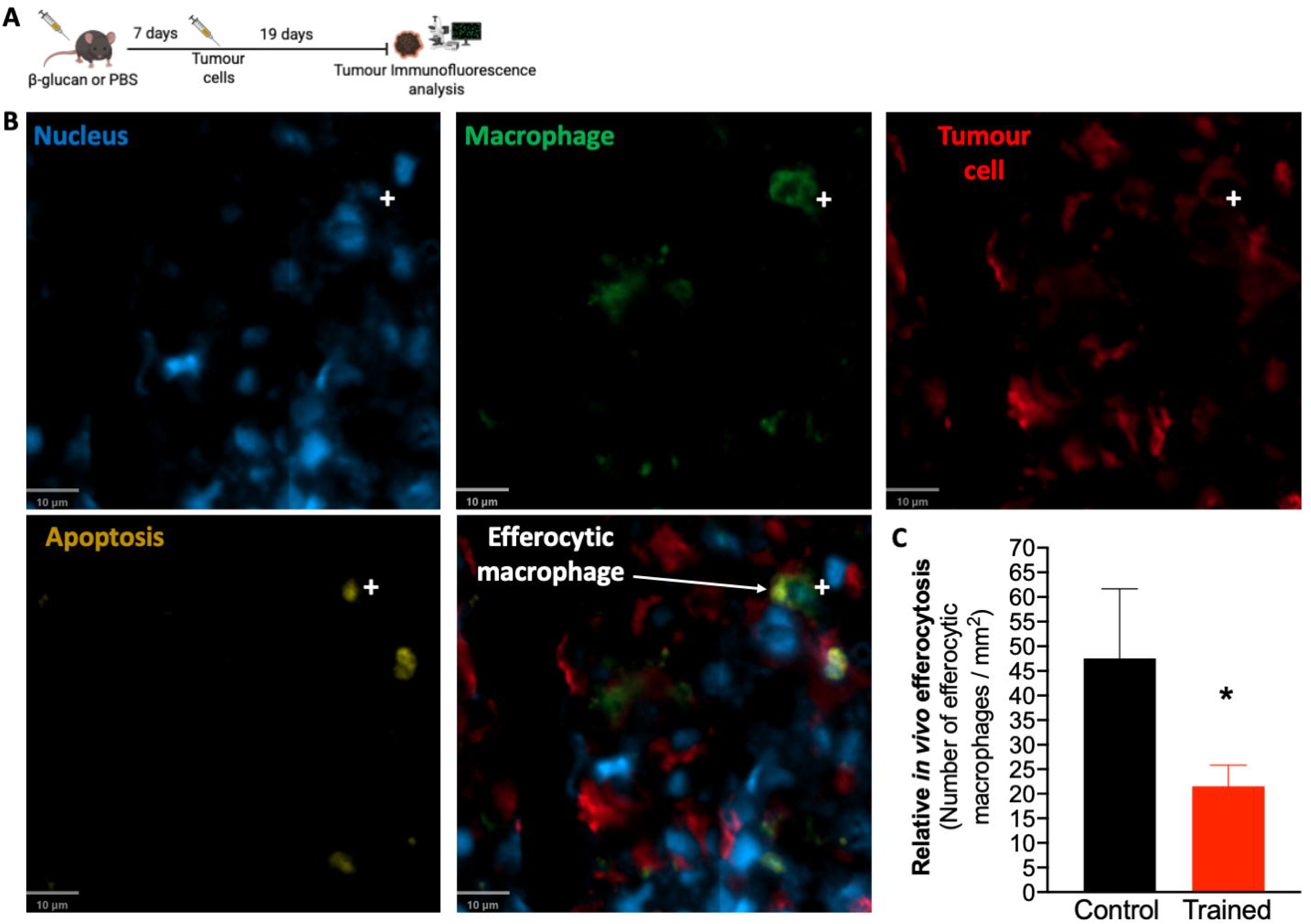
Induction of trained innate immunity inhibits macrophage efferocytosis of tumour cells *in vivo*. (A) Experimental scheme is shown (created in Biorender). (B, C) WT mice were injected with a single i.p. injection of β-glucan or PBS and 7 days thereafter, mice were subcutaneously inoculated with B16-F10 melanoma cells. Tumour tissues were collected for histology 19 days after B16-F10 injection. (B) Representative images of immunofluorescence microscopy of B16-F10 tumour cryosections stained with anti-F4/80, anti-TA99 and anti-cleaved caspase 3 (cCasp3) antibodies (Scale bar: 10 μm). The symbol shows a phagocytic macrophage. (C) Quantification plot of macrophages that have engulfed apoptotic tumour cells, i.e. efferocytic macrophages, (F4/80+ TA99+ cCasp3+) in non-necrotic regions of B16F10 tumours (n = 3 control mice; n = 4 trained mice). The levels of relative *in vivo* efferocytosis are presented as the number of efferocytic macrophages per mm^2^ of non-necrotic tumour tissue. \**P*<0.05, one-tailed Student’s *t*-test. Data are presented as mean ± s.e.m.

**Supplementary Figure 4.**
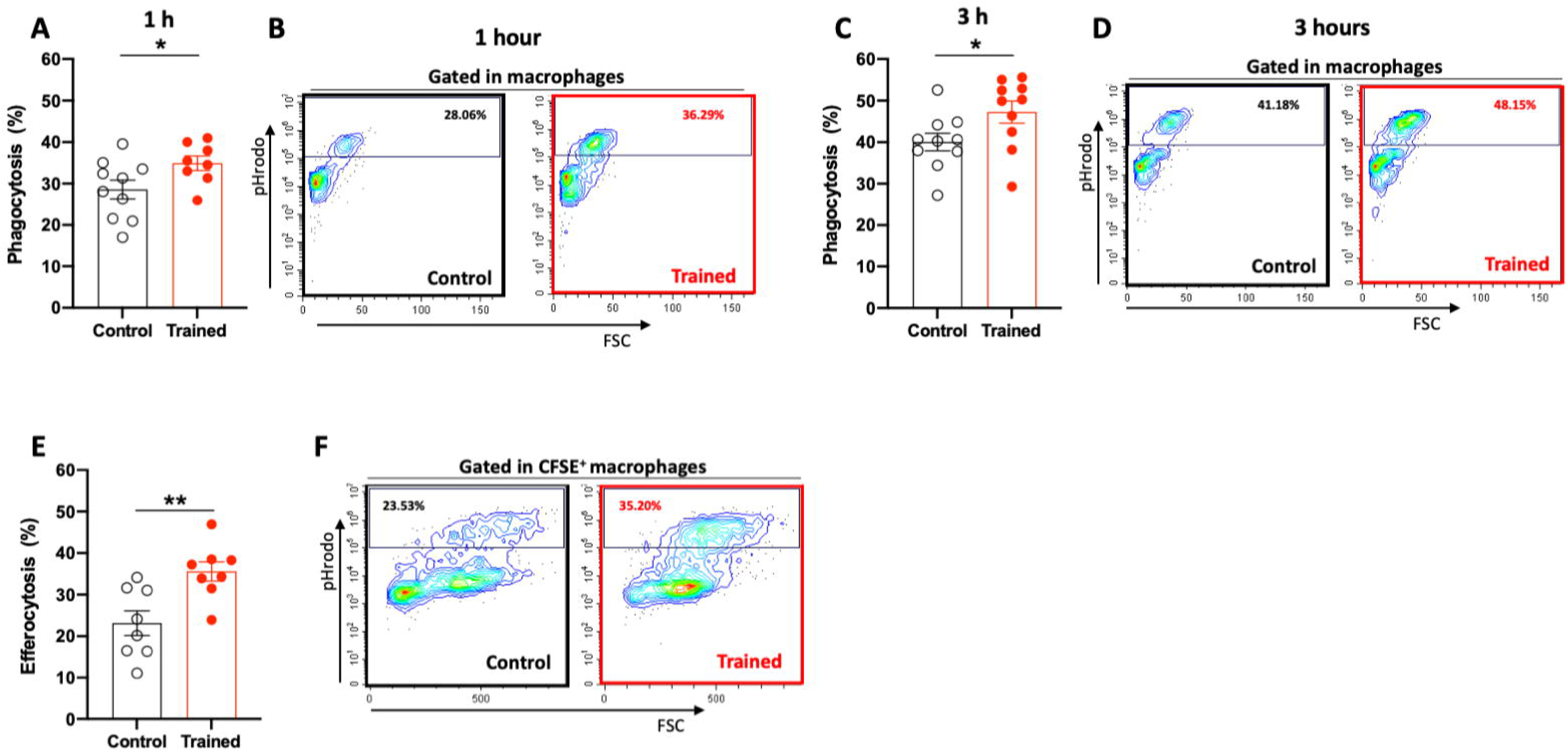
The role of trained innate immunity on macrophage phagocytosis of microbes and apoptotic neutrophils. (A-D) Trained or control macrophages were treated with 50 μg/ml of pHrodo green *E. coli* particles for 1 h and 3 h. Percentage of macrophages that have engulfed *E. coli* particles is shown (A, C: *n*LJ=LJ8-10 separate cell isolations per group). (E, F) Trained or control macrophages, pre-stained with CFSE, were co-cultured with pHrodo red – labelled apoptotic neutrophils for 30 min. Percentage of macrophages that have engulfed apoptotic neutrophils is shown (*n*=8 separate cell isolations per group). (B, D, F) Representative fluorescence-activated cell sorting plots for *E. coli* particle phagocytosis (B, D) and neutrophil efferocytosis (F) are shown. Numbers in outlined areas indicate the percentage of macrophages that is pHrodo^+^ green (B, D) and pHrodo^+^ red (F). \**P*<0.05, \*\**P*<0.01. Two-tailed Student’s *t*-test (A, C, E). Data are presented as mean ± s.e.m. and are pooled from two experiments (A, C) or derived from one experiment (E).

**Supplementary Figure 5.**
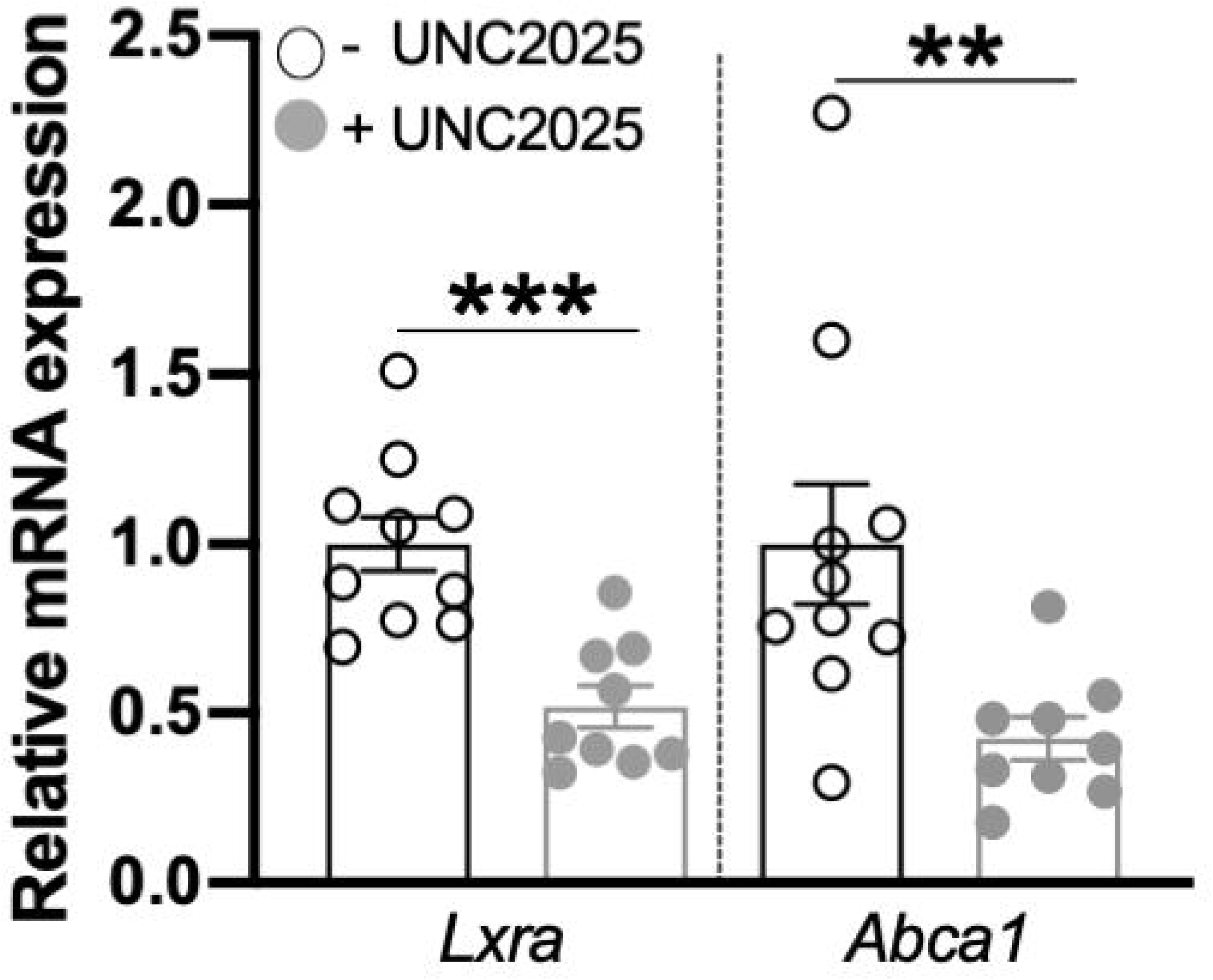
The effect of MerTK blockade on macrophage gene expression. Relative mRNA expression of *Lxra* and *Abca1* in macrophages that were treated or not with 0.1 μΜ of the MerTK inhibitor UNC2025 for 60 minutes. Relative mRNA expression was normalized against 18S rRNA and was set as 1 in the untreated macrophages (n=9-10 separate cell isolations per group). \*\**P*<0.01, \*\*\**P*<0.001. Two-tailed unpaired *t*-test test. Data are presented as mean ± s.e.m. and are pooled from two experiments.

